# One for the road: Bumble bees consume pollen at flowers

**DOI:** 10.1101/2024.06.10.598376

**Authors:** Maggie M. Mayberry, Jacob S. Francis, Jenny K. Burrow, Faith E. Dall, Michelle Bowe, Anne Leonard, Parker Campbell, Avery L. Russell

**Affiliations:** Department of Biology, Missouri State University, Springfield, MO, 65897. USA; Department of Biological Sciences, Florida Atlantic University, Boca Raton FL, 33431, USA; Department of Biology, University of Nevada, Reno, Reno, NV, 89557. USA

**Author notes:** Corresponding author Department of Biology, 910 S John Q Hammons Pkwy, Temple Hall, Missouri State University, Springfield, MO 65897 USA, email of corresponding author.

**Keywords:** Bumble bees, foraging, pollen, pollen foraging, nectar, nutritional ecology

## Abstract

Bees are the primary consumers of pollen in many ecosystems, but pollen consumption by adult bees is rarely studied, leaving a gap in our understanding of the nutritional ecology of collective foraging and pollination biology more generally. For example, while eusocial bees feed upon pollen from colony stores, whether they also consume pollen directly from flowers to meet their own needs or to assess its quality for the broader collective is unknown. We therefore captured wild bumble bee colonies (*B. bimaculatus* and *B. griseocollis*) and tested whether individual workers consumed pollen directly from flowers in a lab-based foraging assay. After confirming presence of floral pollen in worker crops (i.e., consumption at flowers), in a field setting we tested alternative hypotheses for the function of this behavior using information about the composition, abundance, and diversity of pollen found in the crops vs. pollen baskets of three species of pollen– and nectar-foraging bumble bees (*Bombus bimaculatus, B. griseocollis,* and *B. impatiens*). Consistent with the hypothesis that consuming pollen at flowers reflects sampling, total pollen quantity in crops was consistently smaller than in pollen baskets, and basket pollen tended to be a subset of that found in crops. Further, pollen foragers consumed more and different kinds of pollen than nectar foragers. Pollen consumption at flowers is thus unlikely to be purely incidental, or to substantially benefit workers nutritionally. Instead, consuming pollen directly from flowers likely allows foragers to quickly assess pollen quality before collecting it to feed the colony as whole.

**SIGNIFICANCE STATEMENT:** While nectar-collecting bees are classic models for the study of foraging behavior and plant-pollinator interactions, little is known about how bees assess pollen while foraging. Pollen is a critical source of protein and lipids, offered by many plant species alongside or instead of nectar. Though variation in pollen macronutrient content and secondary chemistry affect bee reproductive performance and health, whether and how foragers evaluate pollen quality is not known. We show that foraging worker bumble bees consume pollen at flowers and suggest this behavior may allow them to sample its nutritional quality. This sheds new light on the nutritional basis of plant-pollinator interactions and adds to our understanding of how bees regulate their collection of this critical resource.

## INTRODUCTION

When dietary needs differ within a social group or across life stages, the decisions made by generalist foragers can reflect a complex mix of nutritional priorities (rev. Lihoureau et al. 2018). For example, parents may encounter trade-offs between their own nutritional requirements and those of offspring, such as when oviposition decisions made by adults are influenced by the quality of nectar or pollen that they consume, rather than just the leaf chemistry that their future offspring will encounter (rev. Sheirs et al. 2002; Wackers et al. 2006). Perhaps to mitigate some of these potential conflicts, social insects in particular have developed sophisticated systems of nutrient regulation that allow the diverse dietary requirements of various life stages and castes to be met effectively (Senior et al. 2016 and references within).

In this context, generalist bees are a model system for investigating how individual foraging decisions are shaped by diverse dietary demands (Helm et al. 2017; Lihoreau et al. 2018). Solitary and social bee species must collect floral resources to meet both their own nutritional needs and those of larvae, which sometimes overlap and sometimes differ substantially (Brodschneider and Crailsheim 2010). For example, both adult bees and larvae require a baseline level of protein, carbohydrates and lipids, and diets rich in particular nutrients such as fatty acids are beneficial to both adult and larval performance. Yet worker and larval pollen nutritional needs are also often expected to differ in significant ways (Paoli et al. 2014; Helm et al. 2017; Kraus et al. 2019; Austin et al. 2021). For instance, adult honey bees prioritize carbohydrates over essential amino acids, whereas honey bee larvae require abundant essential amino acids over carbohydrates (Paoli et al. 2014; Helm et al. 2017). These essential macronutrients are unevenly distributed across floral resources, with pollen serving as the primary source of protein, essential lipids, and vitamins (Roulston et al. 2000; Vaudo et al. 2020) for most bee species, and nectar serving as bees’ major source of carbohydrates. To further complicate matters, pollen and nectar produced by different plant species can vary in both macronutrient content and secondary chemistry in ways that are meaningful to both larval and adult performance (e.g., Praz et al. 2008; Stabler et al. 2015; Richardson et al. 2015).

Given that both adult bees and larvae need to consume pollen, it is notable that pollen feeding at flowers by adult foragers is rarely reported. Among at least some solitary bee species, female adult bees do consume pollen directly from flowers to meet their own nutritional needs (Cane 2016; Urban-Mead et al. 2022; Nagano et al. 2023). However, to maintain and develop flight muscles, glandular secretions, and gonads, adults of social species such as honey bees, bumble bees, and stingless bees are largely reported to consume pollen from larval provisions (Roulston and Cane 2000; Smeets and Duchateau 2003; Pech-May et al. 2012; Grund-Mueller et al. 2020; Nagano et al. 2023) rather than at flowers. Consuming pollen directly from flowers, rather than from larval provisions, could have multiple benefits for eusocial species. For instance, eusocial bees are typically highly generalized in diet, and colonies collect pollen from potentially dozens of plant species (Kriesell et al. 2017; Vaudo et al. 2018; Yourstone et al. 2023). Floral pollen consumption could enable adult bees to sample the nutritional quality of pollen before collecting it into their external pollen baskets. Alternately, floral pollen consumption could allow adults to efficiently pursue their own nutritional needs separately from those of the larvae (Paoli et al. 2014; Helm et al. 2017; Tanaka et al. 2019; Kraus et al. 2019; Austin et al. 2021), as they could presumably digest pollen collected in their crop while contributing pollen in their pollen baskets to the colony. Yet whether eusocial worker bees consume pollen directly from flowers, and why they might do so is an open question.

In this study, we addressed two basic questions about the pollen foraging behavior of bumble bees. As a starting point, we asked (Q1) under lab and field settings, whether foragers of three species consume pollen at flowers. Having robustly established that this behavior occurs, we next sought to understand its function (Q2). We used information about the type, quantity, and diversity of pollen we observed in bees’ crops vs. pollen baskets to evaluate predictions of three alternative hypotheses (Figure 1). While we do not consider these hypotheses to be mutually exclusive, we identified *a priori* predictions that would be consistent with each serving as the major explanation for observed patterns in pollen found in crops and baskets.

**Figure 1.**
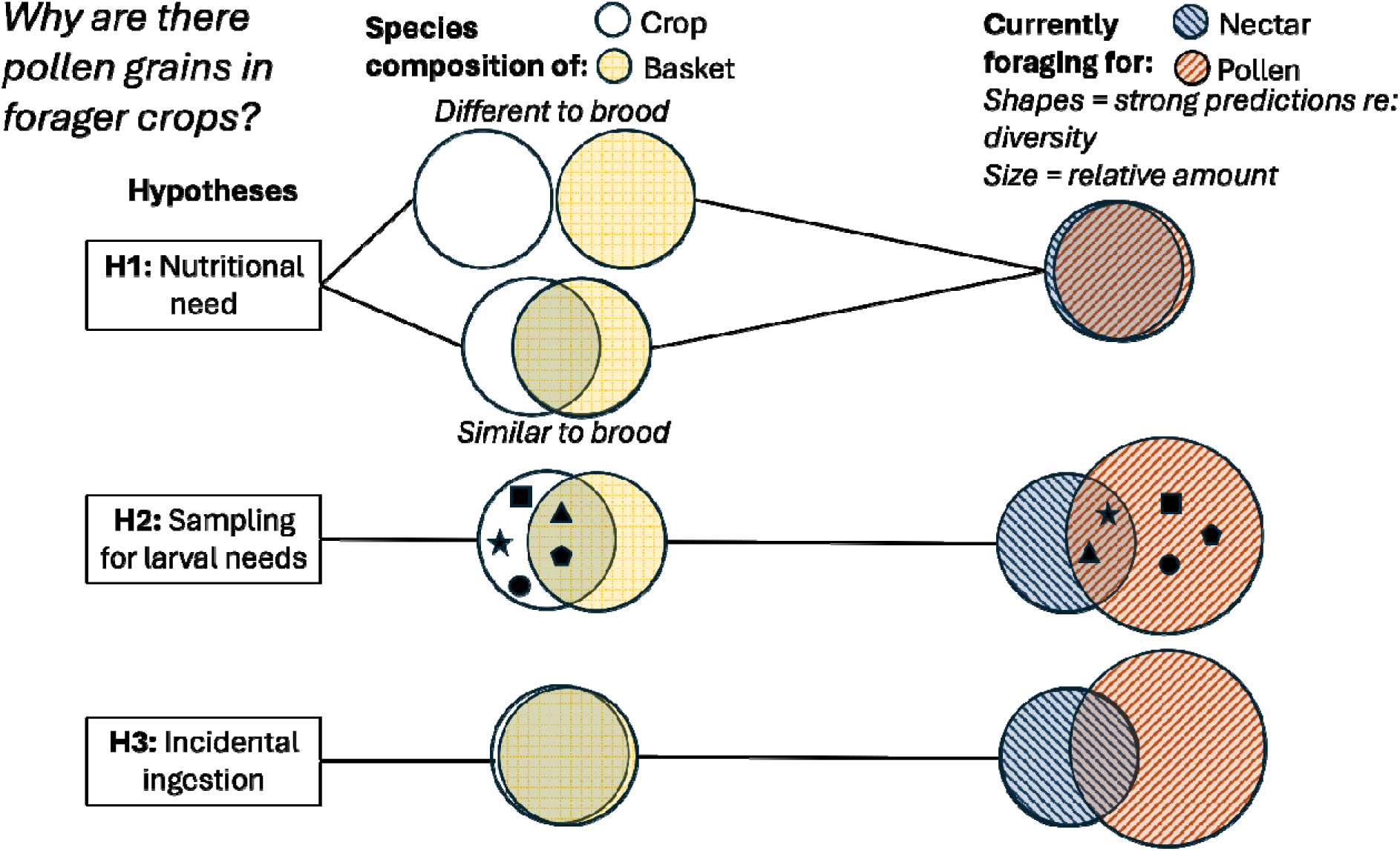
Conceptual diagram illustrating the three hypotheses considered and tested in this study.

First, we considered the hypothesis that workers consume pollen directly from flowers for their own nutrition (Figure 1, H1, top row). If so, we expected to find relatively similar crop pollen composition and amount among nectar and pollen foraging workers, assuming they would seek out preferred pollen types that would be relatively similar despite showing short-term specialization on different foraging tasks (e.g., Russell et al. 2017). We also expected to find some degree of difference in the composition of crop vs. basket pollen (largely destined for larval consumption). We expect little compositional overlap if dietary needs differ between workers and larvae, but it is also possible that compositional differences are minor if quantity (rather than quality) is the major difference in need across life stages.

Second, we also considered the hypothesis (Figure 1, H2, middle row) that floral pollen consumption could be a means by which pollen foragers assess pollen nutritional quality, before collecting it for larval provisions. Like nectar, pollen from different plant species differs in nutritional qualities, including amino acid, lipid, and protein content, and pollen foragers frequently express preferences for a given pollen type (Roulston and Cane 2000; Kriesell et al. 2017; Wood et al. 2019; Vaudo et al. 2020; Ruedenauer et al. 2018, 2020, 2021). Might adult bees also assess pollen quality by consuming it directly from flowers? If so, we expected that among pollen foragers, pollen types in pollen baskets should be a subset of pollen types found in crops. Accordingly, when comparing pollen vs. nectar foraging workers, we also expected pollen foragers to consume more pollen and/or more diverse pollen than nectar foragers.

Finally, we also considered the null expectation (Figure 1, H3, bottom row) that pollen found in forager crops simply reflects incidental ingestion. Pollen foraging can be messy, and while packing pollen into pollen baskets bumble bees regurgitate nectar, potentially exposing their mouthparts to whatever is present in the local environment. Perhaps it is simply difficult to keep pollen grains out of crops, but its consumption has no behavioral or nutritional significance. If so, we expected to find no significant differences in the composition of crop and basket loads among all foragers. By definition, we would expect to find more pollen in the crops of pollen foragers, but no differences in diversity, nor similarity in the pollen composition of nectar and pollen forager crops, because foragers typically visit different plant sources for nectar vs. pollen (e.g., Brian 1957; Somme et al. 2015).

## METHODS

### Laboratory experiment

To test whether bumble bees consume pollen directly from flowers while foraging, we maintained 2 wild-caught colonies (one of *Bombus bimaculatus* and one of *B. griseocollis*) in the lab, following Russell et al. (2017). We allowed bees from each colony to forage freely on 2 M sucrose solution from artificial feeders within enclosed test arenas (LWH: 82 x 60 x 60) set to a 14:10 h light:dark cycle. To ensure that any pollen in the crop was directly consumed while foraging on flowers, colonies were not provided pollen during this short experiment.

In each behavioral trial we assessed whether pollen consumption occurred directly from flowers being visited, rather than consumed from colony stores, by mounting 9 fresh *Solanum houstonii* flowers, a species native to Mexico and southern Arizona, in a 3×3 array on the arena wall in a cleaned test arena. This plant species offers only pollen as a reward (flowers are nectarless). Flowers were grown in a university greenhouse with supplemental halogen lights to extend day length to a 14:10 h light: dark cycle, and were fertilized weekly (PlantTone, NPK 5:3:3, Espoma, Millville, New Jersey, USA). From the foraging arena, a single female worker bee was gently captured from the nectar feeder using a 40 dram vial (Bioquip Products, Inc.) and released in the test arena. During a single trial, we allowed a bee to forage on the 9 flowers for a maximum of 40 visits (mean visits ± SE: 33 ± 3). After the trial, the flowers were discarded and the bee was immediately euthanized and stored in a –20° C freezer for later identification and quantification of pollen in crops and pollen baskets.

We used cleaned fine forceps (Bioquip Products, Inc.) to completely remove the pollen from each pollen basket (‘corbicula’) and store them in 1mL of 70% ethanol in microcentrifuge tubes. We then carefully dissected abdomens to remove the crops, which were stored in microcentrifuge tubes in 70% ethanol at –20° C. We examined the crops, rather than the mid or hind gut, because the crop contents reflect very recent consumption (e.g., pollen passes from the crop to the midgut within 30 minutes in honey bees; Peng et al. 1986). To permit pollen counting, we used cleaned forceps to break open crops and vortexed them for 30 seconds to release the pollen. We discarded crop fragments and checked that discarded fragments did not contain pollen using a compound light microscope. We condensed crop samples to 100uL via centrifuge and vortexed samples prior to counting and identification.

### Field experiment

In order to examine pollen in the crops and pollen baskets of bees foraging under natural conditions, from June 24 – July 8 2020, we collected female bumble bees visiting flowers in the Waterwise Garden (Springfield, MO: 37.18683704788533, –93.27686340229961). Because the Waterwise Garden has a high density and diversity of flowering plant species (at least 52 flowering species based on our collection) and is surrounded by lawns composed primarily of grasses (e.g., *Cynodon <u>dactylon</u>*, but also clover species) and roads, we assumed that bees within the garden likely foraged almost entirely within it. To identify pollen grains in the collected crops and pollen baskets, we created a physical and digital pollen reference collection for all flowers at the Waterwise Garden that were blooming during our field study. Each flower was photographed and identified to species and we collected anthers from each species into 70% ethanol in separate 1.5mL microcentrifuge tubes. Pollen from each sample was vortexed and mounted on separate reference slides and digitally photographed at 400 × magnification using a compound light microscope (3000-LED Series, Accu-Scope, Inc.).

Bees were collected from 1100 hours to 1600 hours, when we observed the maximum number of worker bees. Pollen and nectar foraging bumble bees were collected directly from flowers into 40 dram vials (Bioquip Products, Inc.), which were temporarily stored in a chilled cooler. We characterized a bee as a pollen forager if she had pollen in her pollen baskets, though ‘pollen foragers’ often drink nectar while foraging (Francis et al. 2016; Russell et al. 2017). Bees were subsequently euthanized in a –20° C freezer for later identification to species (Williams et al. 2014) and analysis of crop and pollen basket contents (as described above).

For all crop and corbiculae samples we counted pollen in three 10 uL aliquots using a hemocytometer (Hausser Scientific, Horsham, PA) at 400 × under the compound light microscope to estimate pollen quantity and composition for the total volume. We simultaneously identified each pollen grain to plant species using our pollen reference collection. If we counted zero grains in these three aliquots we counted grains in all remaining available aliquots. Total estimated pollen counts were rounded to the nearest whole number.

### Data Analyses

All data were analyzed using R v.4.4.0 (R Development Core Team 2024). Broadly, we built generalized linear models to test for differences among forager types and tested for significant effects via likelihood ratio tests. We used simulated residuals to assess whether a given model’s residual errors met assumptions and took a stepwise approach to account for overdispersion, non-normality of residuals, and heteroskedasticity (DHARMa: Hartig 2022). For visualization, we calculated estimated marginal effect sizes and 95% confidence errors (emmeans: Lenth 2012).

### Q1: Do bumble bees consume pollen directly from flowers while foraging?

We compared how much pollen captive and field-collected pollen foragers had in their crops using a negative binomial generalized linear model (GLM) via the lme4 package (Bates et al. 2015), with fixed effects for bee origin (‘field-collected’ vs ‘captive’) and bee species (‘*B. bimaculatus*’, ‘*B. griseocollis*’, ‘*B. impatiens*’). To test if field-collected pollen or nectar foraging bees were more likely to have pollen in their crops, we used a two proportion Z-test in base R.

### Q2: Why do foragers consume floral pollen?

To address H1 (Figure 1), we tested whether field-collected pollen foragers had more pollen in their crops than nectar foragers using a negative binomial GLM, with fixed effects for forager type (‘pollen’ vs ‘nectar’) and bee species (‘*B. bimaculatus*’, ‘*B. griseocollis*’, ‘*B. impatiens*’). In a separate analysis, we used a hurdle modeling approach to test if pollen foraging bees were more likely to have more than one species of pollen in their crops than nectar foraging bees via a binomial GLM. We then tested whether the pollen community was more diverse (richness, Shannon’s and Simpson’s diversity indices) for nectar versus pollen foragers (only for bees with pollen in their crops) using a negative binomial GLM (for richness) or binomial GLM (for Shannon’s and Simpson’s), with fixed effects as above. We added 0.01 to all Shannon’s and Simpson’s values to correct for overdispersion.

To address H2 (Figure 1), we examined whether the composition (Bray-Curtis dissimilarity) of pollen loads in the crops of pollen and nectar foragers differed, using a permutation ANOVA (vegan package; Oksanen et al. 2024), specifying bee species and forager type as fixed effects.

To address H3 (Figure 1), in field-collected bumble bees that were actively foraging for pollen, we tested whether the composition of pollen loads in the corbiculae (pollen baskets) and crop of pollen foragers differed as above, specifying bee species and pollen origin as fixed effects. Additionally, we tested whether pollen species richness differed between crops and corbiculae using a generalized linear mixed effects model (GLMM), with fixed effects for pollen origin (‘crop’ versus ‘pollen basket) and bee species and a random intercept of individual forager (to account for non-independence of crops and corbiculae). Only bees with pollen in their crops were considered for these analyses.

## RESULTS

### Bumble bees consume pollen directly from flowers while foraging

In a lab setting, we found that bees readily collected and consumed pollen directly from flowers: 93% of bumble bees (*N* = 15) from captive field-collected *B. bimaculatus* and *B. griseocollis* colonies consumed pollen directly from fresh *Solanum houstonii* flowers. We also confirmed that this behavior also appeared to occur at our field site, where all three species of bumble bees (*Bombus bimaculatus, B. impatiens, B. griseocollis*) had pollen in their crops, comprising 45% of the 58 flowering plant species that were in bloom. On average, captive pollen foraging bees had 846 pollen grains (± SE: 190) in their crops, which was 5.6 times as much pollen as field-collected pollen foraging bumble bees (Figure 2a; GLM: bee origin effect: χ^2^ = 14.65, *P* < 0.0002; bee species effect: χ^2^ = 4.65, *P* = 0.098). Pooling nectar and pollen foragers across bee species, individual worker bees had on average two types of pollen and 308 pollen grains in their crops (Table 1). Pollen and nectar foragers were equally likely to have pollen in their crops (percent of pollen vs nectar foragers with pollen in their crops: 88% vs 82%, respectively; two proportion Z-test: χ^2^ = 0.214, *P* = 0.644).

**Figure 2.**
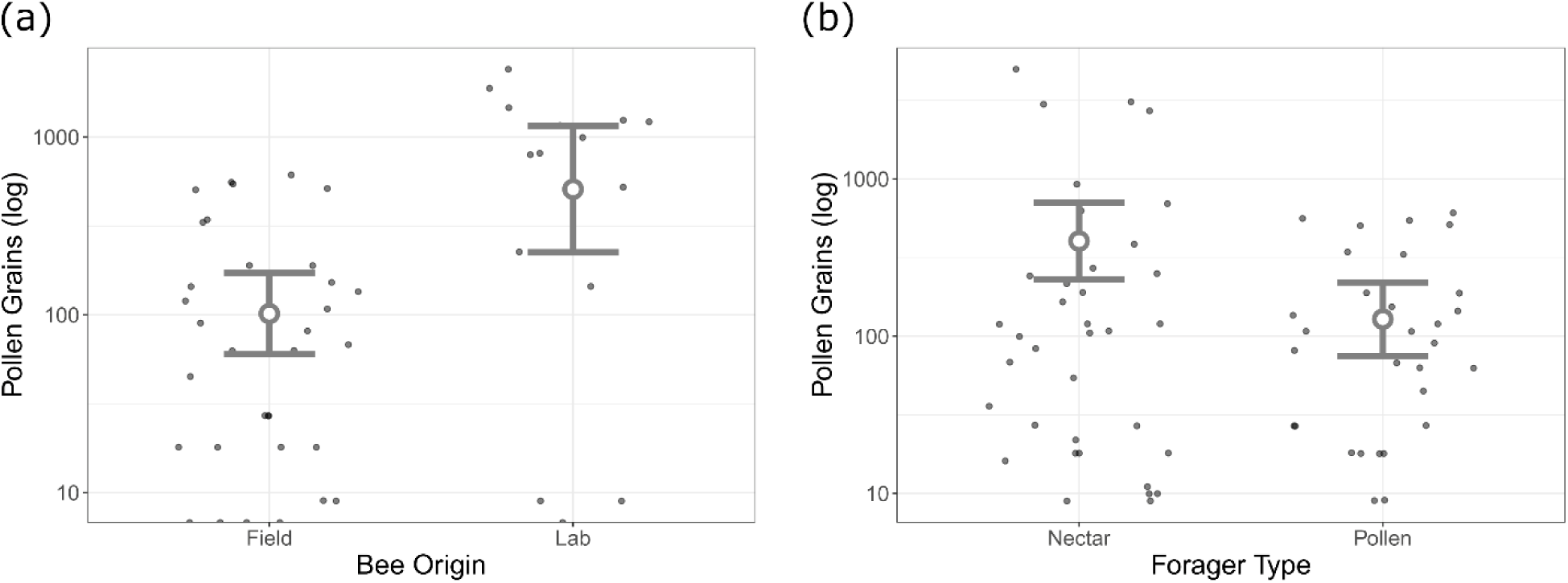
Quantity of pollen grains in the crops of bumble bees. Comparison of **(a)** captive and field-collected pollen foragers and **(b)** field-collected nectar and pollen foragers. For **(a)** *N* = 15 and 34 captive and field-collected pollen foragers, respectively; for **(b)** *N* = 36 and 30 field-collected nectar and pollen foragers, respectively (bees with pollen in crops). Plotted with 95% confidence intervals.

Table 1.

**Table 1.**
|  | Pollen foragers<br>(*N*=34 bees) | Nectar foragers (*N*=44<br>bees) | Pooled foragers<br>(*N*=78 bees) |
| --- | --- | --- | --- |
| Mean ± SE pollen<br>grains in crop | 151 ± 32 | 429 ± 185 | 308 ± 89.3 |
| Mean ± SE pollen<br>types in crop | 2.5 ± 0.34 | 1.75 ± 0.26 | 2 ± 0.21 |
| Diversity (Shannon<br>Index) of pollen in<br>crop | 0.735 | 0.439 | 0.574 |
| Diversity (Simpson<br>Index) of pollen in<br>crop | 0.436 | 0.280 | 0.351 |
| Mean ± SE pollen<br>grains in pollen<br>basket | 1090111 ± 239985 | | |
| Mean ± SE pollen<br>types in pollen<br>basket | 6 ± 0.54 | | |
| Diversity (Shannon<br>Index) of pollen in | 0.659 ± 0.072 | | |
| pollen basket |  |  |  |
| Diversity (Simpson Index) of pollen in pollen basket | $0.334 \pm 0.037$ | | |

### Why do foragers consume floral pollen?

If floral pollen consumption reflects foragers’ personal nutritional needs (Figure 1, H1), then we expected that in our field experiment, we would find similar patterns across nectar and pollen foragers in terms of amount, composition and diversity of crop pollens. In contrast to this, across bee species, nectar foragers consumed 2.8 times as much pollen as pollen foragers (Table 1), a difference that was statistically significant (Figure 2b; GLM: forager type effect: χ^2^_1_ = 10.20, *P* < 0.0014; bee species effect: χ^2^_2_ = 2.47, *P* = 0.291). Despite nectar foragers consuming more pollen grains, pollen foragers were 23% more likely to have more than one species in their crops (Figure 3a; ΔAIC = 2.18, χ^2^_1_ = 4.18, *P* < 0.041) and there was a trend for pollen foragers to have consumed a greater species richness of pollen (Figure 3b; Table 1; GLM: forager type effect: χ^2^_1_ = 2.88, *P* = 0.090; bee species effect: χ^2^_2_ = 0.17, *P* = 0.919). Additionally, pollen foragers consumed pollen from a significantly greater diversity of plant species than nectar foragers (Figure 3c; Table 1; GLM: Shannon–Wiener diversity index: forager type effect: χ^2^_1_ = 5.80, *P* < 0.016; bee species effect: χ^2^_2_ = 0.13, *P* = 0.935). However, when measured via Simpson’s diversity index, pollen in nectar forager crops was more diverse, suggesting that pollen in pollen forager crops is dominated by a few species (Figure 3d; GLMs: forager type effect: χ^2^_1_ = 4.77, *P* < 0.029; bee species effect: χ^2^_2_ = 0.59, *P* = 0.746). Finally, the composition (Bray-Curtis dissimilarity) of pollen within crops of pollen and nectar foragers was significantly different (PERMANOVA: *F* = 6.58, *P* < 0.011).

**Figure 3.**
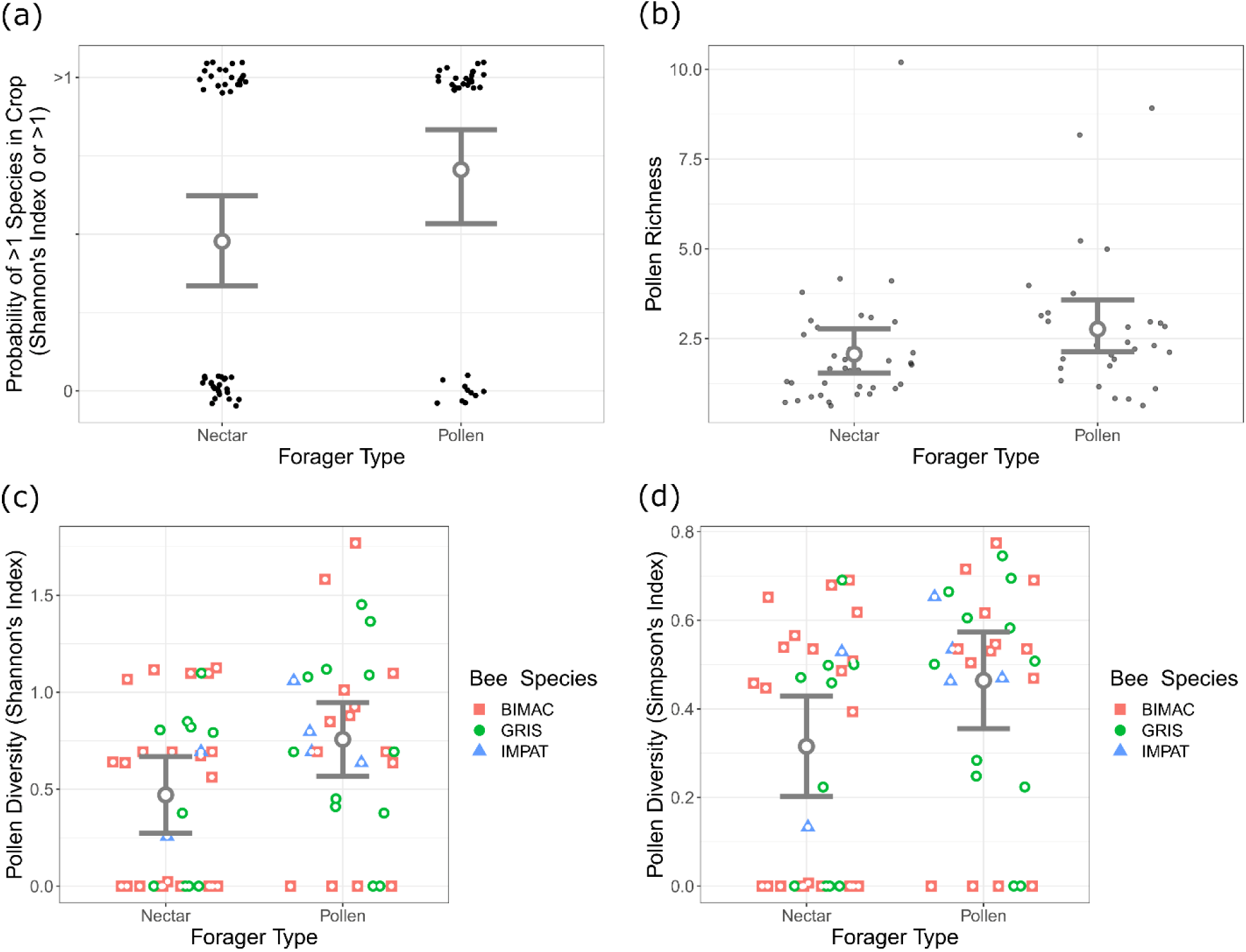
Patterns of pollen richness and diversity in the crops of field-collected nectar and pollen foraging bumble bees. **(a)** Probability that a bee consumed just one species, **(b)** species richness of consumed pollen, and **(c,d)** diversity of consumed pollen measured via **(c)** Shannon’s or **(d)** Simpsons’s diversity index. **(a)** *N* = 44 and 34 nectar and pollen foragers, respectively; **(b-d)** *N* = 36 and 30 nectar and pollen foragers, respectively (bees with pollen in crops). BIMAC = *B. bimaculatus*; GRIS = *B. griseocollis*; IMPAT = *B. impatiens*. Plotted with 95% confidence intervals.

If consuming pollen directly at flowers represents sampling of quality (Figure 1, H2), then we expected to find that pollen basket contents would be a subset of a wider diversity of pollen grains found in the crop (i.e., rejected pollen sources would not be represented in the pollen baskets). Consistent with this, compositional similarity of pollen in corbiculae was greater than for pollen in crops, suggesting that pollen collected within the pollen baskets was overall a subset of pollen consumed by pollen foragers (Figure 4a).

**Figure 4.**
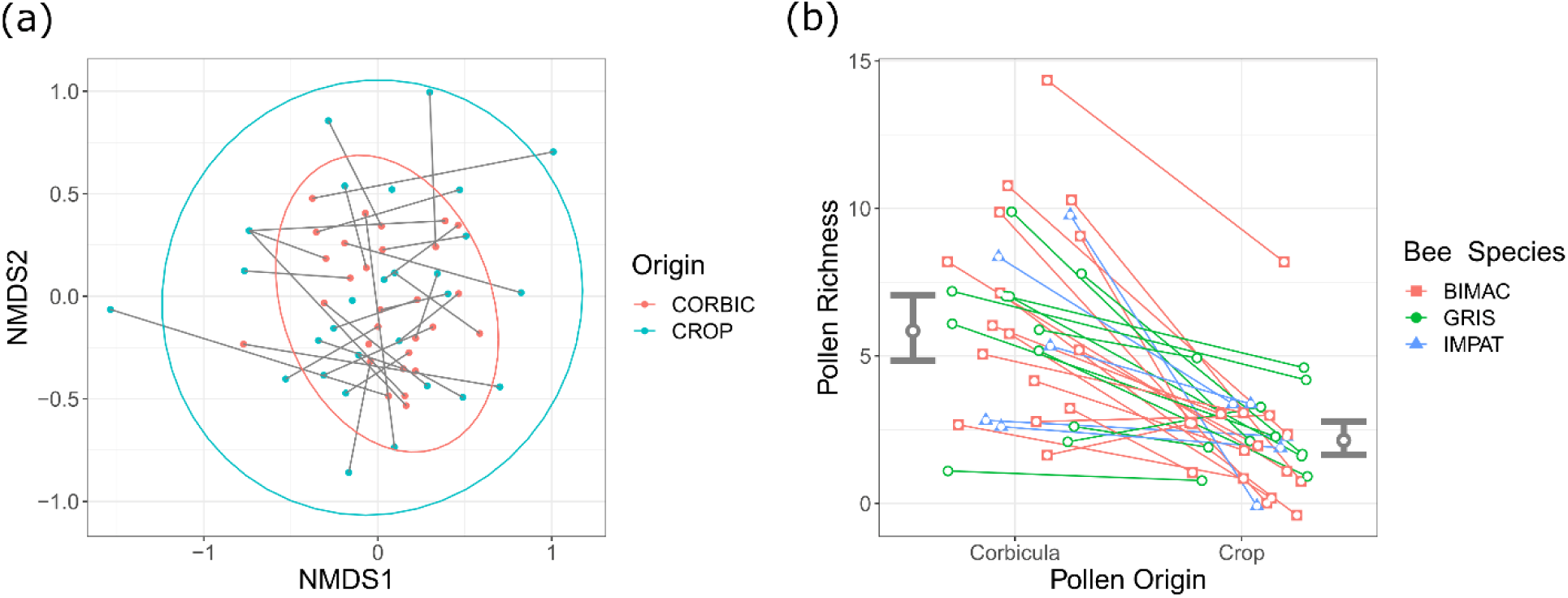
Patterns of pollen richness and composition in the pollen baskets and crops of field-collected pollen foragers. **(a)** Nonmetric multidimensional scaling (NMDS) plot of pollen specie composition and **(b)** species richness of pollen in pollen baskets (‘CORBIC’) and crops; lines join crops and pollen baskets of the same bee. *N* = 30 foragers (only bees with pollen in crops). BIMAC = *B. bimaculatus*; GRIS = *B. griseocollis*; IMPAT = *B. impatiens*. **(a)** Plotted with 95% ellipsoids or **(b)** confidence intervals.

If floral pollen consumption simply reflects incidental ingestion (Figure 1, H3), we expected there would be no distinction between the crop and baskets in terms of composition or diversity. While the composition of pollen within pollen baskets and crops of pollen foragers did not differ overall (PERMANOVA: *F* = 0.052, *P* = 0.816), we found that richness of pollen grains in pollen forager pollen baskets was 2.4 times that of their crops (Table 1), a difference that was statistically significant (Figure 4b; GLMM: forager type effect: χ^2^_1_ = 60.38, *P* < 0.00001; bee species effect: χ^2^_2_ = 0.20, *P* = 0.904).

## DISCUSSION

Our study elucidates patterns of direct pollen consumption from flowers by worker bumble bees and potential functions of this poorly understood behavior. We found that regardless of bumble bee species, nearly all field-collected nectar and pollen foragers had pollen in their crops. Given that crop contents likely constitute evidence of very recent feeding (Peng et al. 1986; Cane 2016), on its own this result suggests the pollen had been consumed during foraging trips. In addition, in the lab, a majority of pollen foragers from wild-caught bumble bee colonies were observed consuming pollen directly from flowers. Together, these results constitute strong evidence that bumble bee workers typically consume pollen directly from flowers as they forage and that this pattern may be widespread among bumble bee species. Furthermore, bumble bees likely do not consume pollen directly from flowers primarily for their own nutrition, given that the composition of pollen in crops of nectar and pollen foragers differed and the seemingly nutritionally trivial amount of pollen found in the crops of lab and field-collected bees (less than 0.03% of what bees collected into their pollen baskets). Instead, we propose that consuming small amounts of pollen directly from flowers is primarily a means by which foragers assess pollen quality. Consistent with this hypothesis, across all pollen foragers, the diversity of pollen collected into the pollen baskets appears to be a subset of the diversity of pollen in the crops.

The quality of both nectar and pollen rewards varies widely and foraging bees are expected to attend to these differences (Roulston et al. 2000; Vaudo et al. 2020; Nicolson 2022). Chemoreceptors on the antennae, proboscis, and even tarsi are frequently used by adult bees while foraging on flowers to assess nectar quality even without ingesting the nectar (Wykes 1952; De Brito Sanchez et al. 2014). As a result, workers can assess which flowers are most rewarding while foraging and even learn to associate specific flower cues with the nectar reward (Giurfa 2007; Kaczorowski et al. 2012; Schiestl and Johnson 2013; Foster et al. 2014). Such associative learning is in fact thought to be fundamental to bee foraging success (Menzel 2012; Chittka and Thomson 2005). However, because pollen nutrients are locked within the tough pollen exine (Roulston and Cane 2000), how adult bees learn to associate floral cues with pollen rewards (e.g., Muth et al. 2016; Russell et al. 2016) and whether bees can assess pollen quality at all while foraging (e.g. Ruedenauer et al. 2018, 2020, 2021) has remained frustratingly unclear (Nicholls and Hempel de Ibarra 2016). Assuming that bees can access and assess pollen nutrients soon after consuming the pollen, our results indicate bumble bees should be able to make decisions related to pollen quality while foraging on flowers.

Why then did bumble bees foraging for nectar also consume small quantities of pollen directly from flowers? One possibility is that workers continuously (re)assess pollen quality opportunistically (see Ruedenauer et al. 2016) even while visiting flowers primarily for nectar, and only switch to pollen collection once they discover an acceptable pollen type. Alternatively or additionally, these bees might be assessing pollen quality while foraging for nectar in order to inform their pollen collecting decisions on subsequent foraging bouts. Little is known about why individual bumble bees switch between foraging for pollen versus nectar, but while some workers readily switch between food types in a single bout, others specialize on only nectar or pollen on a given foraging bout (Goulson 1999; Goulson et al. 2002; Russell et al. 2017). Additional experiments will be required to determine whether and when the quality of consumed pollen influences nectar foraging bees to switch to pollen foraging.

Conversely, pollen consumption on flowers by nectar foragers may have little to do with informing the worker’s future pollen foraging decisions. Instead, consuming pollen continuously could have pharmacological benefits. Pollen often contains high levels of secondary metabolites (Palmer-Young 2019), some of which are implicated in improving memory (e.g., Wright et al. 2015; Jones and Agrawal 2022) and control of parasites (e.g., Baracchi et al. 2015; Richardson et al. 2015; Giacomini et al. 2018). Thus, nectar or even pollen foragers might consume pollen while foraging to continuously self-medicate (a revised and testable version of our H1). Alternatively, or in addition, direct pollen consumption may instead be at least in part an accidental byproduct of consuming nectar (Figure 1, H3). Nectar often contains pollen in quantities similar to what we recovered from bee crops (Herrera et al. 2017). Given that nectar foragers had, unexpectedly, on average nearly three times as much pollen in their crops as pollen foragers, consumption of pollen-contaminated nectar might at least partially explain this pattern.

Interestingly, workers from wild-caught bumble bee colonies in the lab consumed on average more than five times as much pollen from flowers as did workers captured in the field. Assuming that pollen consumption is primarily a means to assess pollen quality, perhaps field bees consumed less pollen, because they were already familiar with the available plant species, unlike the captive workers, which had been provided a novel plant species. On the other hand, the diet of captive colonies was protein deficient relative to the field bees, since we did not supplement the colonies with pollen. Thus, perhaps workers of captive colonies were consuming pollen directly from flowers to supplement their nutrition. Accordingly, honey bee workers that are initially pollen deprived, when later offered pollen, compensate by eating more pollen (Brodschneider et al. 2022).

In conclusion, our study opens the door for studying the potential function(s) of pollen consumption on flowers for eusocial bees. While our results do not preclude the possibility that pollen in the crops of field-collected worker bees could have in part been consumed from colony stores, at a minimum we demonstrate that bumble bees will consume pollen from diverse plant species while foraging, even when that pollen is concealed within the flower (e.g., such as within the tube-like poricidal anthers of the *Solanum* flowers we provided in the lab). At our field site, of the 58 plant species in bloom, bees overall collected and consumed pollen from 34 and 26 species, respectively. However, individual workers were much more specialized and on average consumed pollen from only two plant species and collected pollen from only six plant species (or two, if excluding non-host pollen; i.e., pollen that comprised less than 5% of a given sample; Cane and Sipes 2006). These results are consistent with prior work, which has demonstrated that while an individual bumble bee colony often has a generalized pollen diet, individual workers on a given foraging bout are typically quite specialized (e.g., Mayer et al. 2012; Somme et al. 2015; Kriesell et al. 2017; Yourstone et al. 2023). Future work will be required to determine whether consumed pollen types are nutritionally inferior to those ultimately collected into the pollen baskets and how consuming pollen directly from flowers affects the health and behavior of eusocial foraging bees, including beyond bumble bees.

## Supporting information

Data and R script used in analyses

## ACKNOWLEDGEMENTS

We are grateful to Jeff Maddox for finding and donating wild-caught bumble bee colonies, Abilene Mosher for greenhouse care, and Russell lab members for discussion. We acknowledge this work was performed on unceded traditional territory of the Kiikaapoi, Sioux, and Osage.

This work was partially supported by NSF Award #2109460 (to JSF).

## ETHICS APPROVAL

All bumble bee experimentation was carried out in accordance with the legal and ethical standards of the USA.

## CONSENT TO PARTICIPATE

Not applicable

## CONSENT FOR PUBLICATION

Not applicable

## DATA ACCESSSIBILITY

The datasets supporting this article are available as electronic supplementary material.

## AUTHOR CONTRIBUTIONS

MMM and ALR conceived and designed the study. MMM, JKB, FED, PC, and ALR collected the data and performed the lab experiment. JSF, ALR, and MMM analyzed the data. MMM, ALR, and AL wrote the original draft of the manuscript; other authors provided editorial advice.

