## Supplementary material for "One for the road: Bumble bees consume pollen at flowers": Data and R script used in analyses: Pollen_Consumption_Analysis_6-7-2024.R

```
#
# Last edited: 2/22/23

# 2/22/23: Reran Shannon's and Simpson's diversity indexes with removal of bees that consumed 0 pollen grains
        #  Found minor differences in mean values for indexes, but both still failed zero inflation test
        #  Will need to try negative binomial glm instead
# 2/27/23: Spent an ungodly amount of time trying to learn how to edit violin plots and then I finally realized I could reuse Avery's code and then it only took like 20 minutes. BUT I know a LOT about making violin plots now, so that's cool. 
        #  Line 303 has exportable plot for PF vs NF Pollen Richness
        #  Will work on formatting other plots next
# 3/1/23:  Created figures to put in ms. Main issue is still formatting. The lines are too thin and the text is too small and the whole thing is grainy. Decided this was a future issue though and instead focused on getting figures into ms draft.
        #  Will likely need to rework the "Density" or "Abundance" figure for NF vs. PF bc someting seems weird 
        #  Will need to pull values to write figures legends next

# 5/24/24: All analyses and figures completed

# 6/7/24: Fixed NZ figure, all species color and shape coordinated

####################

# Load Required Packages####
library(plyr)
library(vegan)
library(ggplot2)
library(lme4) #utils::install.packages("lme4")
library(DHARMa)
library(MASS)
library(emmeans)
library(multcomp)
library(reshape2)
library(car)
library(DHARMa)
library(asbio)
library(viridis)

# Get citations from packages
citation("vegan")


rm(list=ls()) #note this will clear your workspace. Its a good thing to do if you 
              #have a bunch of things floating around, but if you have big objects
              #you want to save, save them before running this

##  set working directory, you shouldn't have to do this 
#   if you open the code from the folder supplied
setwd(dirname(rstudioapi::getActiveDocumentContext()$path))

# Set plot directory 
PLOTDIR <- dirname(rstudioapi::getActiveDocumentContext()$path)


# --LAB BEES-- ####

# Examine whether there is more pollen in the crops of field bees vs lab bees

## read in the data  ####
LabVsFieldBees<-read.csv("Processed_lab-vs-field-bees_corbicula-and-crop.csv",stringsAsFactors = T)

### Plot Density ###

LabVsFieldBees$Density  <- LabVsFieldBees$Pollen_Count
Density2 <- ggplot(LabVsFieldBees, aes(x=Bee_Origin, y= Density))
Density_Plot2 <- Density2 +
  geom_point(size=1.5,position=position_jitter(width = .3), alpha=.5) +
  scale_x_discrete("Bee Origin",labels=c("Field","Lab"))+
  scale_y_log10("Pollen Grains (log)") +
  theme_bw() + theme(text=element_text(size=18))
Density_Plot2

# A negative binomial model should account for the overdispersion
DensMod5 <- glm.nb(data=LabVsFieldBees, Density ~ Bee_Origin + Bee_Species)

# Test Model Assumptions
simResid_DensMod5 <- simulateResiduals(fittedModel = DensMod5)
plot(simResid_DensMod5, form=LabVsFieldBees$Bee_Origin) # looks good


# Lets do a Likelihood Ratio Test for significance
drop1(DensMod5, test = "Chisq") # Difference by bee origin!

# Density ~ Bee_Origin + Bee_Species
#           Df Deviance    AIC     LRT  Pr(>Chi)    
# <none>           60.346 627.99                      
# Bee_Origin   1   74.994 640.64 14.6472 0.0001296 ***
# Bee_Species  2   64.993 628.64  4.6462 0.0979683 .

modmeansDensity5<- cld(emmeans(DensMod5,specs = ~Bee_Origin))

DensSummary5 <- with(modmeansDensity5, data.frame(Bee_Origin=Bee_Origin,
                                                Density = exp(emmean),
                                                upper = exp(asymp.UCL),
                                                lower = exp(asymp.LCL))
)

DensXOriginBee <- Density_Plot2 + geom_errorbar(data=DensSummary5, aes(ymin=lower, ymax=upper), width=.3, size=2, color=rgb(.5,.5,.5))+
  geom_point(data=DensSummary5, size=6, col=rgb(.5,.5,.5))+
  geom_point(data=DensSummary5, size=3, col="white")

DensXOriginBee

######


# Load in files
FieldBees<-read.csv("Processed_field-bees_corbicula-and-crop.csv",stringsAsFactors = T)
FieldBees_PFCorb <- read.csv("BPC_PF Corbic_2-16.csv")
FieldBees_NFCrop <- read.csv("BPC_NFCrop.csv")
FieldBees_PFCrop <- read.csv("BPC_PFCrop.csv")
FieldBeesCrops <- FieldBees[FieldBees$Origin=="CROP",]  # lets only examine Crops at this point...

# Examine the data 
str(FieldBees)

# Replace 0s with NAs
FieldBees[,-c(1,2,3,4,5,6)][is.na(FieldBees[,-c(1,2,3,4,5,6)])] <- 0 # this bit fo code replaces the NAs with  0s in the plant species columns
plantSpecies <- colnames(FieldBees[,-c(1,2,3,4,5,6)])
str(FieldBees)

FieldBees_PFCorb[,-c(1,2,3,4,5,6)][is.na(FieldBees_PFCorb[,-c(1,2,3,4,5,6)])] <- 0 #
FieldBees_PFCrop[,-c(1,2,3,4,5,6)][is.na(FieldBees_PFCrop[,-c(1,2,3,4,5,6)])] <- 0
FieldBees_NFCrop[,-c(1,2,3,4,5,6)][is.na(FieldBees_NFCrop[,-c(1,2,3,4,5,6)])] <- 0 


# --FIELD BEES-- ####

NZ_FieldBees <- FieldBees[FieldBees$Tot_Pollen_Type!=0,]

## Proportion of NF and PF bees with pollen in crops

# Test for significant difference

croppol_props <- prop.test(x = c(36, 30), n = c(44, 34))
croppol_props
# Not significant X-squared = 0.21389, df = 1, p-value = 0.6437


## Question 1:  Do pollen foragers carry a greater diversity of pollen in their crops than nectar foragers? ####

  # Don't use this because not all of these bees consumed or collected pollen

# Shannon's Index
FieldBees$Shannon  <- apply(FieldBees[,-c(1,2,3,4,5,6)],1,FUN = function(x) diversity(x,"shannon"))
FieldBees_PFCorb$Shannon <- apply(FieldBees_PFCorb[,-c(1,2,3,4,5,6)],1,FUN = function(x) diversity(x,"shannon"))
FieldBees_PFCrop$Shannon <- apply(FieldBees_PFCrop[,-c(1,2,3,4,5,6)],1,FUN = function(x) diversity(x,"shannon"))
FieldBees_NFCrop$Shannon <- apply(FieldBees_NFCrop[,-c(1,2,3,4,5,6)],1,FUN = function(x) diversity(x,"shannon"))

summary(FieldBees$Shannon)        # x = 0.537
summary(FieldBees_PFCorb$Shannon) # x = 0.659
summary(FieldBees_PFCrop$Shannon) # x = 0.649
summary(FieldBees_NFCrop$Shannon) # x = 0.359

# Simpson's Index
FieldBees$Simpson  <- apply(FieldBees[,-c(1,2,3,4,5,6)],1,FUN = function(x) diversity(x,"simpson"))
FieldBees_PFCorb$Simpson  <- apply(FieldBees_PFCorb[,-c(1,2,3,4,5,6)],1,FUN = function(x) diversity(x,"simpson"))
FieldBees_PFCrop$Simpson  <- apply(FieldBees_PFCrop[,-c(1,2,3,4,5,6)],1,FUN = function(x) diversity(x,"simpson"))
FieldBees_NFCrop$Simpson  <- apply(FieldBees_NFCrop[,-c(1,2,3,4,5,6)],1,FUN = function(x) diversity(x,"simpson"))

summary(FieldBees$Simpson)        # x = 0.311
summary(FieldBees_PFCorb$Simpson) # x = 0.334
summary(FieldBees_PFCrop$Simpson) # x = 0.503
summary(FieldBees_NFCrop$Simpson) # x = 0.411

## GLM Models for Shannon & Simpson Indexes

# Shannon GLM 
    FieldBees$ShannonNZ <- ifelse(FieldBees$Shannon>0,1,0)
    ZeroModel <- glm(data=FieldBees[FieldBees$Origin=="CROP",], ShannonNZ ~ Forager_Type , family = "binomial")
    drop1(ZeroModel,test="Chisq")

   # ShannonNZ ~ Forager_Type
   #              Df Deviance    AIC    LRT Pr(>Chi)  
   # <none>            102.10 106.10                  
   # Forager_Type  1   106.28 108.28 4.1772  0.04097 *    
        
    plotdata  <- FieldBees[FieldBees$Origin=="CROP",]
    
    NZProbs <- data.frame(emmeans(ZeroModel, specs="Forager_Type",type="response"))
    
    NZ_Plot  <-  ggplot(plotdata, aes(x=Forager_Type, y=ShannonNZ))+
      geom_point(position=position_jitter(height = .05, width =.1))+
      geom_point(data=NZProbs, aes(y=prob), size=5)+
      theme_bw() + theme(text=element_text(size=18)) +
      geom_errorbar(data=NZProbs, aes(y=prob, ymin=asymp.LCL,ymax=asymp.UCL), width=.3, size=2, color=rgb(.5,.5,.5))+
      geom_point(data=NZProbs, aes(y=prob), size=6, col=rgb(.5,.5,.5))+
      geom_point(data=NZProbs, aes(y=prob), size=3, col="white")+
      scale_x_discrete("Forager Type",labels=c("Nectar","Pollen"))+
      scale_y_continuous("Probability of >1 Species in Crop\n (Shannon's Index 0 or >1)", breaks=c(0,.5,1), labels=c("0","",">1"))
    NZ_Plot
    

    NZModel <- lm(data=FieldBees[FieldBees$Origin=="CROP"&FieldBees$ShannonNZ>0,], Shannon~Forager_Type)
    drop1(NZModel,test="Chisq")  
    simResid_ShanDiv <- simulateResiduals(fittedModel = NZModel)
    plot(simResid_ShanDiv,form=FieldBees[FieldBees$Origin=="CROP"&FieldBees$ShannonNZ>0,]$Forager_Type) # difference in variance b/t pol and nec
    plot(simResid_ShanDiv, form=FieldBees$Bee_Species)
    
    # Simpson GLM 
    FieldBees$SimpsonNZ <- ifelse(FieldBees$Simpson>0,1,0)
    ZeroModel2 <- glm(data=FieldBees[FieldBees$Origin=="CROP",], SimpsonNZ ~ Forager_Type , family = "binomial")
    drop1(ZeroModel2,test="Chisq")
    
    plotdata2  <- FieldBees[FieldBees$Origin=="CROP",]
    
    NZProbs2 <- data.frame(emmeans(ZeroModel2, specs="Forager_Type",type="response"))
    
    ggplot(plotdata2, aes(x=Forager_Type, y=SimpsonNZ))+
      geom_point(position=position_jitter(.01))+
      geom_point(data=NZProbs2, aes(y=prob), size=5)+
      theme_bw() + theme(text=element_text(size=18)) +
      geom_errorbar(data=NZProbs2, aes(y=prob, ymin=asymp.LCL,ymax=asymp.UCL), width=.2)+
      geom_point(data=NZProbs2, aes(y=prob), size=3, color="white")
    
    NZModel <- lm(data=FieldBees[FieldBees$Origin=="CROP"&FieldBees$SimpsonNZ>0,], Simpson~Forager_Type+Bee_Species)
    drop1(NZModel,test="Chisq")  
    simResid_ShanDiv <- simulateResiduals(fittedModel = NZModel)
    plot(simResid_ShanDiv,form=FieldBees[FieldBees$Origin=="CROP"&FieldBees$SimpsonNZ>0,]$Forager_Type) # difference in variance b/t pol and nec
    plot(simResid_ShanDiv, form=FieldBees$Bee_Species)    
   
    
hist(FieldBees[FieldBees$Origin=="CROP",]$Tot_Pollen_Types)
    

    ShanDiv <- lmer(data=FieldBees, Shannon ~ Forager_Type + Bee_Species + Origin + (1|BeeID)) #TALK TO MAGGIE ABOUT THIS!
    summary(ShanDiv)
    drop1(ZeroModel,test="Chisq")
      # Response: Shannon
      # LR Chisq Df Pr(>Chisq)  
      # Forager_Type   6.0693  1    0.01376 *
      # Bee_Species    3.1933  3    0.36277  
    
    # Check Model Assumptions
    simResid_ShanDiv <- simulateResiduals(fittedModel = ShanDiv)
    plot(simResid_ShanDiv, form=FieldBees$Forager_Type) # difference in variance b/t pol and nec
    plot(simResid_ShanDiv, form=FieldBees$Bee_Species)
    
    testZeroInflation(simResid_ShanDiv) # Failed 
    plot(hist(FieldBees$Shannon))  #talk about this with maggie!
    testDispersion(simResid_ShanDiv)
    # Failed zero inflation test, trying negative binomial
    
    ShanDiv2 <- glm.nb(data=FieldBees, Shannon ~ Forager_Type + Bee_Species)
    warnings(ShanDiv2) # Ask Jake about this # negative binomial is just for counts!
    Anova(ShanDiv2)
    # No significant difference in Shannon's Index between PF vs. NF or among bee species
                    #LR Chisq Df Pr(>Chisq)  
      #Forager_Type   2.9335  1    0.08676 .
      #Bee_Species    3.1469  3    0.36953 
    # Will try rerunning this with 0's ommited from dataset
    
# Simpson GLM
    SimpDiv <- lmer(data=FieldBees, Simpson ~ Forager_Type + Bee_Species + Origin + (1|BeeID)) # talk about this!
    summary(SimpDiv)
    Anova(SimpDiv) 
    
    simResid_SimpDiv <- simulateResiduals(fittedModel = SimpDiv)
    plot(simResid_SimpDiv, form=FieldBees$Forager_Type) # difference in variance b/t pol and nec
    plot(simResid_SimpDiv, form=FieldBees$Bee_Species)
    plot(simResid_SimpDiv, form=FieldBees$Origin)
    
    testZeroInflation(simResid_SimpDiv) # Failed 
    testDispersion(simResid_SimpDiv)
    
    SimpDiv2 <- glm.nb(data=FieldBees, Simpson ~ Forager_Type + Bee_Species)
    warnings(SimpDiv2) # Ask Jake about this 
    Anova(SimpDiv2)
    # No significant difference in Simpson's Index between PF vs. NF or among bee species
                    # LR Chisq Df Pr(>Chisq)
      # Forager_Type  0.30511  1     0.5807
      # Bee_Species   0.20503  3     0.9768
    # Will try rerunning this with 0's ommited from dataset

## Diversity Indexes -- With 0's ommitted

# Load in new files

    #FieldBees_PFCorb <- read.csv("BPC_PF Corbic_2-16.csv")      # No removal necessary (No 0s)
    FieldBees_NFCrop_No0s <- read.csv("BPC_NFCrop_No0s.csv")     # 8 bees removed
    FieldBees_PFCrop_No0s <- read.csv("BPC_PFCrop_No0s.csv")     # 4 bees removed
    
      # GLM File
    FieldBees_No0s <- read.csv("Processed_field-bees_corbicula-and-crop_0s-removed_2-22.csv")
    FieldBeesCrops_No0s <- FieldBees_No0s[FieldBees_No0s$Origin=="CROP",] # pulling crop data only
    
    # Change NAs to 0s
    FieldBees_NFCrop_No0s[,-c(1,2,3,4,5,6)][is.na(FieldBees_NFCrop_No0s[,-c(1,2,3,4,5,6)])] <- 0
    FieldBees_PFCrop_No0s[,-c(1,2,3,4,5,6)][is.na(FieldBees_PFCrop_No0s[,-c(1,2,3,4,5,6)])] <- 0 
    FieldBeesCrops_No0s[,-c(1,2,3,4,5,6)][is.na(FieldBeesCrops_No0s[,-c(1,2,3,4,5,6)])] <- 0 
    
# Shannon
      FieldBees_NFCrop_No0s$Shannon <- apply(FieldBees_NFCrop_No0s[,-c(1,2,3,4,5,6)],1,FUN = function(x) diversity(x,"shannon"))
      FieldBees_PFCrop_No0s$Shannon <- apply(FieldBees_PFCrop_No0s[,-c(1,2,3,4,5,6)],1,FUN = function(x) diversity(x,"shannon"))
      FieldBeesCrops_No0s$Shannon <- apply(FieldBeesCrops_No0s[,-c(1,2,3,4,5,6)],1,FUN = function(x) diversity(x,"shannon"))
      
      summary(FieldBees_NFCrop_No0s$Shannon) # x = 0.439 (was x = 0.3594) 
      summary(FieldBees_PFCrop_No0s$Shannon) # x = 0.735 (was x = 0.6485)
      summary(FieldBeesCrops_No0s$Shannon)   # x = 0.574 
    
    # Run GLM Model to compare Shannon Indexes
    ShanDiv_No0s <- glm(data=FieldBeesCrops_No0s, Shannon+0.01 ~ Forager_Type + Bee_Species, family="gaussian")
    
    summary(ShanDiv_No0s)
    Anova(ShanDiv_No0s) 
        #              LR Chisq Df Pr(>Chisq)  
        # Forager_Type   5.7967  1    0.01606 *
        # Bee_Species    0.1337  2    0.93535 
    
    # Lets do a Likelihood Ratio Test for significance
    drop1(ShanDiv_No0s, test = "Chisq") # Difference by forager type!
    
    # Test Model Assumptions
      simResid_ShanDiv_No0s <- simulateResiduals(fittedModel = ShanDiv_No0s)
      plot(simResid_ShanDiv_No0s, form=FieldBeesCrops_No0s$Forager_Type) # no overdisperson
      plot(simResid_ShanDiv_No0s, form=FieldBeesCrops_No0s$Bee_Species)  # no overdisperson
    
      testZeroInflation(simResid_ShanDiv_No0s) # When adding 0.01, it works
      testDispersion(simResid_ShanDiv_No0s)

# Simpson 
      FieldBees_NFCrop_No0s$Simpson  <- apply(FieldBees_NFCrop_No0s[,-c(1,2,3,4,5,6)],1,FUN = function(x) diversity(x,"simpson"))
      FieldBees_PFCrop_No0s$Simpson  <- apply(FieldBees_PFCrop_No0s[,-c(1,2,3,4,5,6)],1,FUN = function(x) diversity(x,"simpson"))
      FieldBeesCrops_No0s$Simpson <- apply(FieldBeesCrops_No0s[,-c(1,2,3,4,5,6)],1,FUN = function(x) diversity(x,"simpson"))
      
      summary(FieldBees_NFCrop_No0s$Simpson) # x = 0.280 (was x = 0.4110)
      summary(FieldBees_PFCrop_No0s$Simpson) # x = 0.436 (was x = 0.5027)
      summary(FieldBeesCrops_No0s$Simpson)   # x = 0.351 (was x = 0.4201)
      
    SimpDiv_No0s <- glm(data=FieldBeesCrops_No0s, Simpson + 0.01 ~ Forager_Type + Bee_Species, family="gaussian")
    
    summary(SimpDiv_No0s)
    Anova(SimpDiv_No0s) 
        #              LR Chisq Df Pr(>Chisq)  
        # Forager_Type   4.7659  1    0.02903 *
        # Bee_Species    0.5857  2    0.74612

    # Lets do a Likelihood Ratio Test for significance
    drop1(SimpDiv_No0s, test = "Chisq") # Difference by forager type!
        
    # Test Model Assumptions
      simResid_SimpDiv_No0s <- simulateResiduals(fittedModel = SimpDiv_No0s)
      plot(simResid_SimpDiv_No0s, form=FieldBeesCrops_No0s$Forager_Type) # no overdisperson
      plot(simResid_SimpDiv_No0s, form=FieldBeesCrops_No0s$Bee_Species)  # no overdisperson
      
      head(FieldBeesCrops_No0s)
      
      testZeroInflation(simResid_SimpDiv_No0s)  # When adding 0.01, it works
      testDispersion(simResid_SimpDiv_No0s)
      


### Plot Richness ####
FieldBeesCrops_No0s$Richness <- FieldBeesCrops_No0s$Tot_Pollen_Types

Richness <- ggplot(FieldBeesCrops_No0s[FieldBeesCrops_No0s$Origin=="CROP",], aes(x=Forager_Type, y= Richness))
Richness_Plot <- Richness +
     geom_point(size=1.5,position=position_jitter(width = .3), alpha=.5) +
     scale_x_discrete("Forager Type",labels=c("Nectar","Pollen"))+
     scale_y_continuous("Pollen Richness") +
     theme_bw() + theme(text=element_text(size=18))
Richness_Plot


### Plot Diversity ###

## Pollen Diversity (Shannon's Index) crops NF vs. PF by Bee Species
Shannon <- ggplot(FieldBeesCrops_No0s, aes(x=Forager_Type, y= Shannon, color=Bee_Species))
Shannon_Plot <- Shannon +  
   geom_point(size=4, position=position_jitter(width = .3, seed=42), aes(shape=Bee_Species)) +
   geom_point(size=1.5,position=position_jitter(width = .3, seed=42), color="White") +
   scale_shape_manual(values=c(15,16,17)) +
   theme_bw() + theme(text=element_text(size=18)) +
   scale_x_discrete("Forager Type",labels=c("Nectar","Pollen")) +
   scale_y_continuous("Pollen Diversity (Shannon's Index)")
Shannon_Plot

modmeansShan <- cld(emmeans(ShanDiv_No0s,specs = ~Forager_Type))

CropShanSummary <- with(modmeansShan, data.frame(Forager_Type=Forager_Type,
                                                 Shannon = emmean,
                                                 upper = upper.CL,
                                                 lower = lower.CL)
)

ShanXForager <- Shannon_Plot + geom_errorbar(data=CropShanSummary, aes(ymin=lower, ymax=upper), width=.3, size=2, color=rgb(.5,.5,.5))+
  geom_point(data=CropShanSummary, size=6, col=rgb(.5,.5,.5))+
  geom_point(data=CropShanSummary, size=3, col="white") + theme(text=element_text(size=18))

ShanXForager

    
## Pollen Diversity (Simpson's Index) crops NF vs. PF by Bee Species
Simpson <- ggplot(FieldBeesCrops_No0s, aes(x=Forager_Type, y= Simpson, color=Bee_Species)) 
Simpson_Plot <- Simpson +  
   geom_point(size=4, position=position_jitter(width = .3, seed=42), aes(shape=Bee_Species)) +
   geom_point(size=1.5,position=position_jitter(width = .3, seed=42), color="White") +
   scale_shape_manual(values=c(15,16,17)) +
   theme_bw() + theme(text=element_text(size=18)) +
   scale_x_discrete("Forager Type",labels=c("Nectar","Pollen")) +
   scale_y_continuous("Pollen Diversity (Simpson's Index)") 
Simpson_Plot

modmeansSimp <- cld(emmeans(SimpDiv_No0s,specs = ~Forager_Type))

CropSimpSummary <- with(modmeansSimp, data.frame(Forager_Type=Forager_Type,
                                                 Simpson = emmean,
                                                 upper = upper.CL,
                                                 lower = lower.CL)
)

SimpXForager <- Simpson_Plot + geom_errorbar(data=CropSimpSummary, aes(ymin=lower, ymax=upper), width=.3, size=2, color=rgb(.5,.5,.5))+
  geom_point(data=CropSimpSummary, size=6, col=rgb(.5,.5,.5))+
  geom_point(data=CropSimpSummary, size=3, col="white") + theme(text=element_text(size=18))

SimpXForager


### Build  some models of richness ####

# there is a pretty good reason not to run a straight linear model (count data)
# this code runs a poisson model then checks residuals (simulated). It shows that
# there is some significant overdispersion... 
# if you want to understand the residuals check out the DHARMa package vignette (https://cran.r-project.org/web/packages/DHARMa/vignettes/DHARMa.html)


### USE THIS FOR RICHNESS & EVENNESS 

# lets try  GLM - it looks like that model is overdispersed
DivMod2 <- glm(data=FieldBeesCrops_No0s, Richness ~ Forager_Type + Bee_Species, family="poisson")
summary(DivMod2)

simResid_DivMod2 <- simulateResiduals(fittedModel = DivMod2)
plot(simResid_DivMod2, form=FieldBeesCrops_No0s$Forager_Type) # difference in variance b/t pol and nec
plot(simResid_DivMod2, form=FieldBeesCrops_No0s$Bee_Species)

testZeroInflation(simResid_DivMod2)
testDispersion(simResid_DivMod2)

# A negative binomial model should account for the overdispersion
DivMod3 <- glm.nb(data=FieldBeesCrops_No0s, Richness ~ Forager_Type + Bee_Species)

simResid_DivMod3 <- simulateResiduals(fittedModel = DivMod3)
plot(simResid_DivMod3, form=FieldBeesCrops_No0s$Forager_Type) # looks goodd

# Lets do a Likelihood Ratio Test for significance
drop1(DivMod3, test = "Chisq") # Difference by forager type!

# Figure 1c results
# Richness ~ Forager_Type + Bee_Species
#               Df Deviance    AIC    LRT Pr(>Chi)  
# <none>            52.639 243.76                  
# Forager_Type  1   56.569 244.88 2.87708  0.08985 .
# Bee_Species   2   53.861 240.18 0.16842  0.91924

modmeans<- cld(emmeans(DivMod3,specs = ~Forager_Type))

RichSummary <- with(modmeans, data.frame(Forager_Type=Forager_Type,
                       Richness = exp(emmean),
                       upper = exp(asymp.UCL),
                       lower = exp(asymp.LCL))
)
  
RichXForager <- Richness_Plot + geom_errorbar(data=RichSummary, aes(ymin=lower, ymax=upper), width=.3, size=2, color=rgb(.5,.5,.5))+
                geom_point(data=RichSummary, size=6, col=rgb(.5,.5,.5))+
                geom_point(data=RichSummary, size=3, col="white") + theme(text=element_text(size=18))
 
RichXForager


mean(FieldBeesCrops_No0s[FieldBeesCrops_No0s$Forager_Type=="POL",]$Richness)
# 2.833333

mean(FieldBeesCrops_No0s[FieldBeesCrops_No0s$Forager_Type=="Nec",]$Richness)
# 2.138889


## How do pollen and nectar foragers crops compare to each other compositionally?####

FieldBeesCrops_No0s[,-c(1,2,3,4,5,6)][is.na(FieldBeesCrops_No0s[,-c(1,2,3,4,5,6)])] <- 0 # this bit fo code replaces the NAs with  0s in the plant species columns
plantSpecies2 <- colnames(FieldBeesCrops_No0s[,-c(1,2,3,4,5,6)])
str(FieldBeesCrops_No0s)

commMat2 <- as.matrix(FieldBeesCrops_No0s[,plantSpecies2])
PermTest <- adonis2(commMat2~Forager_Type*Bee_Species, data=FieldBeesCrops_No0s, method="bray", permutations = 100000) # This is a permanova that suggests that there are differences in pollen composition

Cropdist <- vegdist(commMat2,method="bray")
dispersion <- betadisper(Cropdist, group=FieldBeesCrops_No0s$Forager_Type)
permutest(dispersion, permutations=10000)
boxplot(dispersion)
table(FieldBeesCrops_No0s$BeeID)

# Response: Distances
# Df  Sum Sq  Mean Sq      F N.Perm Pr(>F)  
# Groups     1 0.03368 0.033681 6.5823  10000 0.0139 *
# Residuals 64 0.32748 0.005117


### ANSWER #### 
  # There is a difference in crop richness between pollen and nectar foragers


## Question 2: Do pollen foragers carry a greater quantity of pollen in their crops than nectar foragers? ####

### Plot Density ####
FieldBeesCrops_No0s$Density  <- FieldBeesCrops_No0s$Tot_Pollen_Count
Density <- ggplot(FieldBeesCrops_No0s, aes(x=Forager_Type, y= Density))
Density_Plot <- Density +
  geom_point(size=1.5,position=position_jitter(width = .3), alpha=.5) +
  scale_x_discrete("Forager Type",labels=c("Nectar","Pollen"))+
  scale_y_log10("Pollen Grains (log)") +
  theme_bw() + theme(text=element_text(size=18))
Density_Plot


### Build Models of Density ####

DensMod2 <- glm(data=FieldBeesCrops_No0s, Density ~ Forager_Type + Bee_Species, family="poisson")
summary(DensMod2)

simResid_DensMod2 <- simulateResiduals(fittedModel = DensMod2)
plot(simResid_DensMod2, form=FieldBeesCrops_No0s$Forager_Type) # difference in variance b/t pol and nec
plot(simResid_DensMod2, form=FieldBeesCrops_No0s$Bee_Species)


# A negative binomial model should account for the overdispersion
DensMod3 <- glm.nb(data=FieldBeesCrops_No0s, Density ~ Forager_Type + Bee_Species)


simResid_DensMod3 <- simulateResiduals(fittedModel = DensMod3)
plot(simResid_DensMod3, form=FieldBeesCrops_No0s$Forager_Type) # looks goodd


# Lets do a Likelihood Ratio Test for significance
drop1(DensMod3, test = "Chisq") # Difference by forager type!

# Figure 1a analysis
# Density ~ Forager_Type + Bee_Species
#               Df Deviance    AIC    LRT Pr(>Chi)   
# <none>            81.933 873.73                   
# Forager_Type  1   92.474 882.46 10.20 0.001405 **
# Bee_Species   2   84.745 872.73  2.47 0.290831

modmeansDensity<- cld(emmeans(DensMod3,specs = ~Forager_Type))

DensSummary <- with(modmeansDensity, data.frame(Forager_Type=Forager_Type,
                                         Density = exp(emmean),
                                         upper = exp(asymp.UCL),
                                         lower = exp(asymp.LCL))
)

DensXForager <- Density_Plot + geom_errorbar(data=DensSummary, aes(ymin=lower, ymax=upper), width=.3, size=2, color=rgb(.5,.5,.5))+
                    geom_point(data=DensSummary, size=6, col=rgb(.5,.5,.5))+
                      geom_point(data=DensSummary, size=3, col="white")

DensXForager

###ANSWER####
#Nectar foragers have significantly more pollen (269 grains) in their crops than pollen
#forager (81 grains).


##Question 3: Is the diversity of pollen types greater in the crop versus in the pollen basket?####

# Start by looking only at pollen foragers
FieldBees$Richness <- FieldBees$Tot_Pollen_Types
FieldBees$Density <- FieldBees$Tot_Pollen_Count

FieldBeesPolForagers <- FieldBees[FieldBees$Forager_Type=="POL",
                                  c("BeeID",
                                    "Bee_Species",
                                    "Origin",
                                    "Richness",
                                    "Density",
                                    "Shannon",
                                    "Simpson")]


#for now lets drop the non paired bees
nonpairs <- read.csv("Non matching data for field bees 1-31-2022.csv", header = F)


BadBees <- nonpairs[grep(pattern = "[0-9]", x=nonpairs$V1, perl=T),1]
FieldBeesPolForagers$BeeID %in% BadBees # they are in there!
FieldBeesPolForagers <- FieldBeesPolForagers[!FieldBeesPolForagers$BeeID %in% BadBees,] # drop em

FieldBeesPolSorted <- arrange(FieldBeesPolForagers,BeeID,Origin)  # we have to sort to handle the point jittering


### Plot Richness by Location - Crop vs. Corb ####
## Pollen Richness PF Crop vs. Corb (plot directly comparing individual bees)
    PolForagers <- ggplot(FieldBeesPolSorted, aes(x=Origin,y=Richness))
    LocationPlot <- PolForagers + 
      geom_line(position=position_jitter(.3,seed=42), aes(color=Bee_Species, group=BeeID))+
      geom_point(size=3,position=position_jitter(.3, seed=42), aes(color=Bee_Species, shape=Bee_Species))+
      scale_shape_manual(values=c(15,16,17)) +
      geom_point(size=1.5, position=position_jitter(.3, seed=42), color="White")+
      scale_x_discrete("Pollen Origin",labels=c("Corbicula","Crop"))+
      scale_y_continuous("Pollen Richness")+
      theme_bw() + theme(text=element_text(size=18))
    LocationPlot

    
    
### Model Richness ####

LocationMod1 <- glmer(data=FieldBeesPolSorted, Richness~Origin + Bee_Species + (1|BeeID),family=poisson)

simResid_LocationMod1 <- simulateResiduals(fittedModel = LocationMod1)
plot(simResid_LocationMod1)
plot(simResid_LocationMod1, form = FieldBeesPolSorted$Origin) # looksgood

drop1(LocationMod1, test="Chisq")

mms <- emmeans(LocationMod1, specs = ~Origin)
presummary <- cld(mms)

# Richness ~ Origin + Bee_Species + (1 | BeeID)
#             npar    AIC    LRT   Pr(Chi)    
# <none>           284.89                     
# Origin         1 343.27 60.381 7.818e-15 ***
# Bee_Species    2 281.10  0.203    0.9035


LocationSummary <- with(presummary, data.frame( Origin= Origin,
                                  Richness = exp(emmean),
                                  upper = exp(asymp.UCL),
                                  lower  = exp(asymp.LCL)
                                  ))

nudge=position_nudge(x=c(.45,-.45)) #for pushing means
totalLocation <- LocationPlot +
  geom_errorbar(data=LocationSummary, size=2, width=.2, color=rgb(.5,.5,.5),position=nudge, aes(ymin=lower,ymax=upper))+
  geom_point(data=LocationSummary, size=4, color=rgb(.5,.5,.5),position=nudge) +
  theme(text=element_text(size=18)) +
  geom_point(data=LocationSummary, size=1.5, color="white", position=nudge)
  

totalLocation # with 95% CI, estimated means based on the generalized linear mixed models


### Shannon ####

    FieldBees_ShanDiv <- glm(data=FieldBeesPolForagers, Shannon + 0.01 ~ Origin + Bee_Species, family="gaussian")

    summary(FieldBees_ShanDiv)
    Anova(FieldBees_ShanDiv) 
    drop1(FieldBees_ShanDiv,test="Chisq")

# Test Model Assumptions
    simResid_FieldBees_ShanDiv <- simulateResiduals(fittedModel = FieldBees_ShanDiv)
    plot(simResid_FieldBees_ShanDiv, form=FieldBeesPolForagers$Origin) # no overdisperson
    plot(simResid_FieldBees_ShanDiv, form=FieldBeesPolForagers$Bee_Species)  # no overdisperson

    head(FieldBeesPolForagers)

    testZeroInflation(simResid_FieldBees_ShanDiv)  # When adding 0.01, it works
    testDispersion(simResid_FieldBees_ShanDiv)

### Simpson ####

    FieldBees_SimpDiv <- glm(data=FieldBeesPolForagers, Simpson + 0.01 ~ Origin + Bee_Species, family="gaussian")

    summary(FieldBees_SimpDiv)
    Anova(FieldBees_SimpDiv) 
    drop1(FieldBees_SimpDiv,test="Chisq")

# Test Model Assumptions
    simResid_FieldBees_SimpDiv <- simulateResiduals(fittedModel = FieldBees_SimpDiv)
    plot(simResid_FieldBees_SimpDiv, form=FieldBeesPolForagers$Origin) # no overdisperson
    plot(simResid_FieldBees_SimpDiv, form=FieldBeesPolForagers$Bee_Species)  # no overdisperson

    head(FieldBeesPolForagers)

    testZeroInflation(simResid_FieldBees_SimpDiv)  # When adding 0.01, it works
    testDispersion(simResid_FieldBees_SimpDiv)     


### Plot Diversity ###
    
    ## Pollen Diversity (Shannon's Index) crop vs. corb
    Shannon <- ggplot(FieldBeesPolSorted, aes(x=Origin, y= Shannon, color=Bee_Species))
    Shannon_Plot2 <- Shannon +  geom_point(size=4, position=position_jitter(width = .3, seed=42)) +
      geom_point(size=1.5,position=position_jitter(width = .3, seed=42), color="White") +
      theme_bw() + theme(text=element_text(size=18)) +
      scale_x_discrete("Pollen Origin",labels=c("Corbicula","Crop"))+
      scale_y_continuous("Pollen Diversity (Shannon's Index)")
    Shannon_Plot2
    
    modmeansShanPol <- cld(emmeans(FieldBees_ShanDiv,specs = ~Origin))

    PolShanSummary <- with(modmeansShanPol, data.frame(Origin=Origin,
                                                 Shannon = emmean,
                                                 upper = upper.CL,
                                                 lower = lower.CL)
    )

    ShanXOrigin <- Shannon_Plot2 + geom_errorbar(data=PolShanSummary, aes(ymin=lower, ymax=upper), width=.3, size=2, color=rgb(.5,.5,.5))+
      geom_point(data=PolShanSummary, size=6, col=rgb(.5,.5,.5))+
      geom_point(data=PolShanSummary, size=3, col="white") + theme(text=element_text(size=18))

    ShanXOrigin

    
    
    # Pollen Diversity (Simpson's Index) crop vs. corb
    Simpson <- ggplot(FieldBeesPolSorted, aes(x=Origin, y= Simpson, color=Bee_Species)) 
    Simpson_Plot2 <- Simpson +  geom_point(size=4, position=position_jitter(width = .3, seed=42)) +
      geom_point(size=1.5,position=position_jitter(width = .3, seed=42), color="White") +
      theme_bw() + theme(text=element_text(size=18)) +
      scale_x_discrete("Pollen Origin",labels=c("Corbicula","Crop"))+
      scale_y_continuous("Pollen Diversity (Simpson's Index)") 
    Simpson_Plot2

    modmeansSimpPol <- cld(emmeans(FieldBees_SimpDiv,specs = ~Origin))
    
    PolSimpSummary <- with(modmeansSimpPol, data.frame(Origin=Origin,
                                                       Simpson = emmean,
                                                       upper = upper.CL,
                                                       lower = lower.CL)
    )
    
    SimpXOrigin <- Simpson_Plot2 + geom_errorbar(data=PolSimpSummary, aes(ymin=lower, ymax=upper), width=.3, size=2, color=rgb(.5,.5,.5))+
      geom_point(data=PolSimpSummary, size=6, col=rgb(.5,.5,.5))+
      geom_point(data=PolSimpSummary, size=3, col="white") + theme(text=element_text(size=18))
    
    SimpXOrigin
    
    

###ANSWER####
#The pollen baskets of pollen foragers are significantly more diverse than their crops

#Gutter - for things not yet finished or not necessary####
FieldBeesSortedCrops <- arrange(FieldBeesCrops,BeeID,Origin) 


#Some Compositional analyses####
## How do pollen foragers crops and corbiculae compare to each other compositionally?####

stackedbarplots <- ggplot(FieldBees[FieldBees$Forager_Type=="POL",], aes(x=Origin,group=BeeID))

Species <- read.csv("Plant_List.csv",header=F)
SpeciesMatrix<-FieldBees[,names(FieldBees) %in%  Species$V1]
SpeciesMatrix$BeeID <-FieldBees$BeeID 
SpeciesMatrix$Forager_Type <- FieldBees$Forager_Type
SpeciesMatrix$Origin <- FieldBees$Origin
SpeciesMatrix$TotPol <- FieldBees$Tot_Pollen_Count
SpeciesMatrix<-SpeciesMatrix[SpeciesMatrix$Forager_Type=="POL",]

Test<- melt(SpeciesMatrix,value.name = "Count")
Test<-Test[Test$variable!="TotPol",]
stacked1 <- ggplot(Test,aes(fill=variable,x=Origin,y=Count))
stackedplot1 <- stacked1+geom_bar(position="fill",stat="Identity")+
  facet_grid(~BeeID,margin=margin(0,0,0,0))+
  theme_bw() + theme(text=element_text(size=18)) +
  theme(strip.text = element_blank(),
        panel.spacing = unit(0,"lines"),
        axis.text.x = element_text(angle=90),
        legend.position = "none")

pdf("Plots/Stacked1.pdf",width=10,height=5) #in inches...
stackedplot1
dev.off()


stacked2 <- ggplot(Test,aes(fill=variable,x=Origin,y=Count))
stackedplot2 <- stacked1+geom_bar(position="fill",stat="Identity")+

  theme_bw() + theme(text=element_text(size=18)) +
  theme(strip.text = element_blank(),
        panel.spacing = unit(0,"lines"),
        axis.text.x = element_text(angle=90))

pdf("Plots/Stacked2.pdf",width=10,height=5) #in inches...
stackedplot2
dev.off()


test1 <-SpeciesMatrix[,names(SpeciesMatrix) %in%  Species$V1]

library(magrittr)
percents<-test1 %<>% mutate(rowsum=rowSums(.)) %>% mutate((./rowsum))

percents$BeeID <- SpeciesMatrix$BeeID
percents$Origin  <- SpeciesMatrix$Origin

library(reshape2)
library(tidyverse)

Test<- melt(percents,value.name = "Count")
Test<-Test[Test$variable!="rowsum",]
Stacked<-ggplot(Test, aes(x=Origin,x=BeeID,y=Count))
Stacked + geom_col()


##NMDS of pollen composition in crops vs corbicula

pollenForagers <- FieldBees[FieldBees$Forager_Type=="POL",]
pollenForagers$Tot_Pollen_Count # NMDS wont work on communities that have no members (e.g. no pollen grains!)
beeswithZero <- pollenForagers$BeeID[pollenForagers$Tot_Pollen_Count==0]
pollenForagerswithBoth <- pollenForagers[!pollenForagers$BeeID%in%beeswithZero,]

commMat <- as.matrix(pollenForagerswithBoth[,plantSpecies])
NMDS<-metaMDS(commMat, k=2, trymax = 1000)
stressplot(NMDS)
plot(NMDS) #not that informative!

NMDSscores <-as.data.frame(scores(NMDS,"sites"))
NMDSspecies <-as.data.frame(scores(NMDS,"species"))
NMDSPlot <- cbind(NMDSscores,pollenForagerswithBoth[,c("BeeID","Bee_Species","Origin")])


#convex hulls... not that useful...
CORBChull <- NMDSPlot[NMDSPlot$Origin=="CORBIC",][chull(NMDSPlot[NMDSPlot$Origin=="CORBIC",c("NMDS1","NMDS2")]),]
CROPChull <- NMDSPlot[NMDSPlot$Origin=="CROP",][chull(NMDSPlot[NMDSPlot$Origin=="CROP",c("NMDS1","NMDS2")]),]
hulls <- rbind(CORBChull,CROPChull)

#elipses
loc.elp <- data.frame() #sets up a data frame before running the function.
veganCovEllipse<-function (cov, center = c(0, 0), scale = 1, npoints = 100) 
{
  theta <- (0:npoints) * 2 * pi/npoints
  Circle <- cbind(cos(theta), sin(theta))
  t(center + scale * t(Circle %*% chol(cov)))
}

for(g in levels(NMDSPlot$Origin)){
  loc.elp <- rbind(loc.elp, cbind(as.data.frame(with(NMDSPlot [NMDSPlot$Origin==g,],
                                                                                   veganCovEllipse(cov.wt(cbind(NMDS1,NMDS2),wt=rep(1/length(NMDS1),length(NMDS1)))$cov,center=c(mean(NMDS1),mean(NMDS2))))) ,Origin=g))
}


NMDSgg <- ggplot(NMDSPlot, aes(x=NMDS1, y=NMDS2))
PrelimNMDS <- NMDSgg + geom_point(aes(color=Origin))+
         geom_line(aes(group=BeeID), color=rgb(.5,.5,.5)) +
         theme_bw() + theme(text=element_text(size=18)) +
        stat_ellipse(aes(color=Origin)) # Draw 95% CI around
     #     geom_polygon(data=hulls, aes(color=Origin, fill=Origin),alpha=.3)
PrelimNMDS


## How do crops and corbiculae of pollen foragers compare to each other compositionally?####

PermTest <- adonis2(commMat~Origin*Bee_Species, data=pollenForagerswithBoth, method="bray", permutations = 100000) # This is a permanova that suggests that there are no differences in pollen composition

Cropdist <- vegdist(commMat,method="bray")
dispersion <- betadisper(Cropdist, group=pollenForagerswithBoth$Origin)
permutest(dispersion, permutations=10000)
boxplot(dispersion)
table(pollenForagerswithBoth$BeeID)

# Response: Distances
#          Df  Sum Sq   Mean Sq      F N.Perm Pr(>F)
# Groups     1 0.00067 0.0006704 0.0519  10000 0.8159
# Residuals 57 0.73592 0.0129109

 
 
 
 ## Export the plots! You can change the size here to make them fit your slides!!!! ####
 
 pdf("Plots/Fig1a_DensityXOriginBee.pdf",width=7,height=5) #in inches...
 DensXOriginBee
 dev.off()

 pdf("Plots/Fig1b_DensityXForagerType.pdf",width=7,height=5) #in inches...
 DensXForager
 dev.off()
 
 pdf("Plots/Fig2a_NZ_Plot.pdf",width=7,height=5) #in inches...
 NZ_Plot
 dev.off()
 
 pdf("Plots/Fig2b_RichnessXForagerType.pdf",width=7,height=5) #in inches...
 RichXForager
 dev.off()
 
 pdf("Plots/Fig2c_Shannon_Plot.pdf",width=7,height=5) #in inches...
 ShanXForager
 dev.off()
 
 pdf("Plots/Fig2d_Simpson_Plot.pdf",width=7,height=5) #in inches...
 SimpXForager
 dev.off()

 pdf("Plots/Fig3a_CropVCorb.pdf",width=7,height=5) #in inches...
 totalLocation
 dev.off()
 
 pdf("Plots/Fig3b_CropVCorbShan.pdf",width=7,height=5) #in inches...
 ShanXOrigin
 dev.off()
 
 pdf("Plots/Fig3c_CropVCorbSimp.pdf",width=7,height=5) #in inches...
 SimpXOrigin
 dev.off()
 
 pdf("Plots/Fig3d_PollenForagersNMDS.pdf",width=7,height=5) #in inches...
 PrelimNMDS
 dev.off()
 
```
